# Meeting the challenges of high-dimensional data analysis in immunology

**DOI:** 10.1101/473215

**Authors:** Subarna Palit, Fabian J. Theis, Christina E. Zielinski

## Abstract

Recent advances in cytometry have radically altered the fate of single-cell proteomics by allowing a more accurate understanding of complex biological systems. Mass cytometry (CyTOF) provides simultaneous single-cell measurements that are crucial to understand cellular heterogeneity and identify novel cellular subsets. High-dimensional CyTOF data were traditionally analyzed by gating on bivariate dot plots, which are not only laborious given the quadratic increase of complexity with dimension but are also biased through manual gating. This review aims to discuss the impact of new analysis techniques for in-depths insights into the dynamics of immune regulation obtained from static snapshot data and to provide tools to immunologists to address the high dimensionality of their single-cell data.

## Curse of Dimensionality in Manual Gating

Since the invention of the first fluorescence-based flow cytometer fifty years ago, immunologists have widely adopted the technology to get a comprehensive understanding of heterogeneity among immune cells, function, cellular differentiation, signaling pathways and biomarker discovery [1]. A variation of flow cytometry, known as cytometry by time-of-flight (CyTOF) or mass cytometry, was developed in 2009, which could query over 50 parameters per cell, in contrast to only limited parameters in conventional flow cytometry. CyTOF utilizes antibodies labelled with rare earth metal isotopes instead of fluorescent dyes and the resulting abundances are detected using a time-of-flight mass spectrometer [2][3]. The preferred high-throughput method for measuring cell surface or intracellular protein abundances depends on certain characteristics that distinguish one technology over the other (**Table 1**).

**Table 1:**
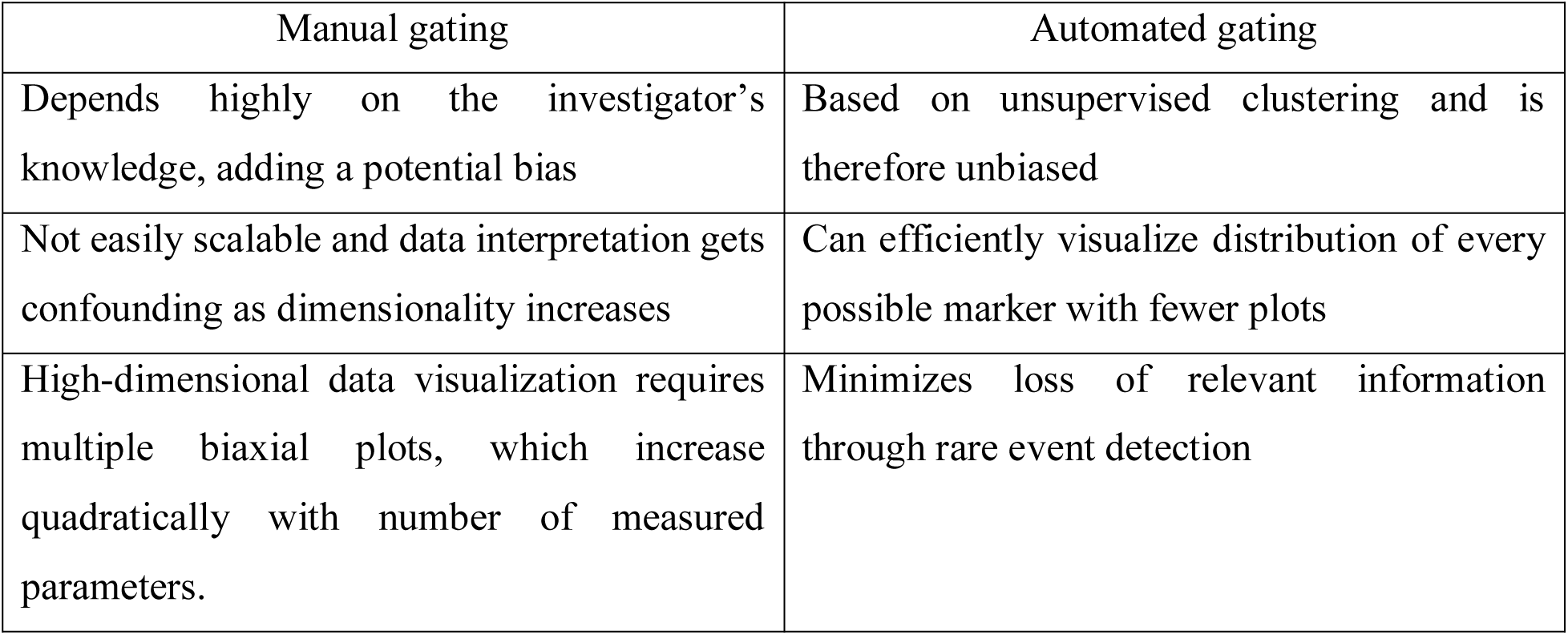
Towards higher dimensions with CyTOF

Analysis of high-dimensional single-cell cytometry data relies on technological advancements and novel analytical methods that can efficiently incorporate the inherent multi-parametric characteristics of such data sets. The most straightforward and traditional, albeit labor-intensive, method for cytometry data analysis is by a process known as “gating”, which uses a series of 2D plots to identify regions of interest in a bivariate scatter plot of single cells [6]. A series of gates drawn in sequence can reveal information about cellular hierarchy and identify subsets of interest from a population. Nevertheless, this approach has several drawbacks when compared to automated strategies (**Table 2**). Data analysis can be handled in one of several ways as new methods are continuously emerging and the shortcomings of manual gating can jeopardize the validity of an experimental finding. A number of automated gating strategies were developed for this purpose. They were designed for cell population identification reproducing expert manual gating and also for sample classification according to some external variables. Several of these methods performed well statistically when compared to manual gating and have been reviewed extensively as part of the first FlowCAP challenge (http://flowcap.flowsite.org/) [7].

**Table 2:**
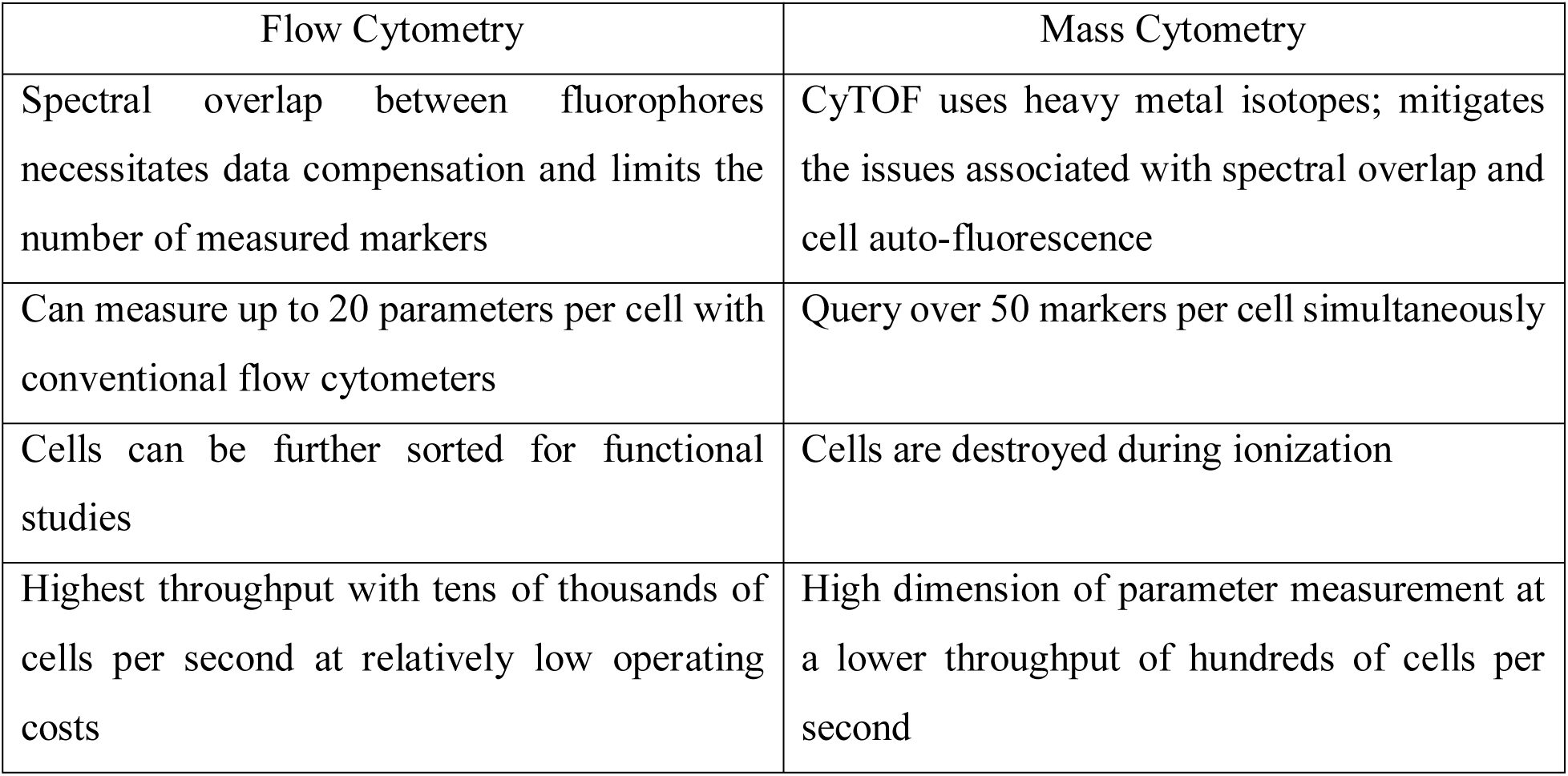
Curse of dimensionality in manual gating

Another rapidly evolving technology, which is taking center stage, is single-cell RNA sequencing (scRNA-seq). It can comprehensively profile the molecular information of individual cells and provides transcriptomics as an additional orthogonal modality, with almost full genomic coverage [5]. Despite potential challenges, rapid computational advancements in single-cell analysis have significantly enabled systematic investigations into cellular heterogeneity, dynamics as well as regulatory mechanisms in an increasing number of tissues.

## Paving the Way for More Complex Analyses

This review aims to discuss newly emerged data analysis tools for downsizing the above-mentioned drawbacks. Analyzing complex high-dimensional single-cell data comes with its own computational challenges and can be reduced to data pre-processing, normalization, dimensionality reduction and clustering followed by cluster biomarker identification (**Figure 1A-C**). Many analysis tools that exist for scRNA-seq data can already be applied to cytometry studies with certain parameters optimized accordingly. To enable transition from experiment to data analysis, many algorithms have already been deployed in the form of interactive visual tools for bench scientists without a need of programming skills. This review delivers a guide for cytometry data analysis by discussing some of the available algorithms including, but not limited to, t-SNE [8], diffusion maps [9], SPADE [10], and FlowSOM [11]. For the first time, we propose two single-cell trajectory inference algorithms, diffusion pseudo-time (DPT) [14] to infer pseudo-temporal ordering of cells according to relative protein abundances from cytometry data, and partition-based graph abstraction (PAGA) [15] for generating network topologies from single-cell data (**Table 3**). Monocle was the first method to quantitatively estimate the progress of acell along some biological or developmental pathway and termed it pseudotime [12]. Another algorithm, known as Wanderlust, uses local topological clustering to align single cells onto a unified developmental path [13]. Originally proposed for the analysis of scRNA-seq, we believe these algorithms hold great potential to uncover the immune system’s cellular composition and differentiation trajectories in a heuristic manner given the growth in dimensions.

**Figure 1.**
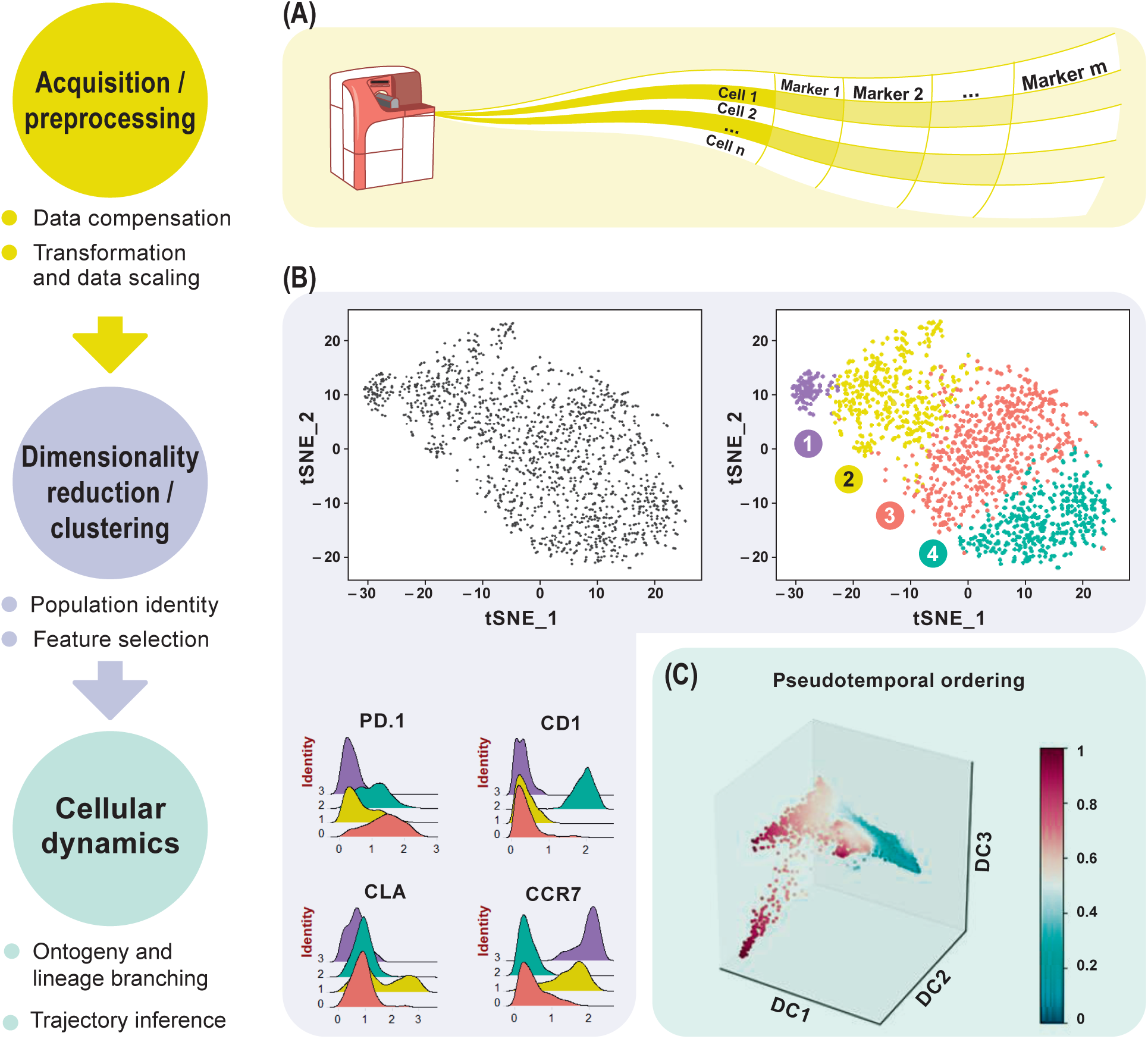
Overview of high-dimensional flow cytometry data analysis. **(A)** FCS files are read as the input data matrix, followed by data compensation (if necessary), transformation, concatenation (in case of multiple input files) and scaling. As an alternative to the standard log transformation method, logicle and arcsinh transformation can be applied. Scaling ensures that every marker in the matrix gets equal importance in further downstream analyses. **(B)** For subsequent analyses, informative markers, such as lineage-specific markers, need to be selected. This can be accomplished via diagnostic plots for marker expression distribution across all events or with prior knowledge. Dimensionality reduction using PCA, tSNE, diffusion maps etc. for visualization follows after feature selection. These plots show similarities between events, measured in an unsupervised manner, and provide information about differential expression between groups based on the intensity measurements. This reveals cellular heterogeneity. **(C)** Detection of developmental trajectories is another common application of these analytical frameworks and provides dynamic information on static snap shot data.

**Table 3:** Overview of some of the established single-cell analysis methods

| Methods | Description | Class |
| --- | --- | --- |
| **PCA** | Cannot account for the smooth nature of single-cell data | Linear dimensionality reduction |
| **t-SNE** | More intuitive representation of high-dimensional data on a lower manifold | Non-linear dimensionality reduction |
| **UMAP** | Scales better and improves global structure of the data compared to t-SNE |  |
| **HSNE** | Scales better than conventional t-SNE |  |
| **Diffusion maps** | Explores continuity through progression of cell differentiation |  |
| **SPADE** | Hierarchical branched tree representation | Clustering methods; single-cell resolution is lost |
| **FlowSOM** | Self-organizing maps trained to detect cellular population |  |
| **Diffusion pseudotime** | Investigates continuous cellular differentiation trajectories from static snapshot single-cell data | Trajectory inference and graph abstraction |
| **Partition-based graph abstraction** | Reconstitutes topological information from complex differentiation single-cell data in the form of cellular maps by applying |  |
|  | strategies of clustering and trajectory inference |  |

## Visualizing Cellular Heterogeneity by Dimensionality Reduction

Flow cytometry represents one of the most powerful and frequently used technologies in the immunologist’s toolbox. Therefore, single-cell resolution has been a hallmark of immunological data acquisition and analysis for a long time since. One of the goals of performing differential analyses of cytometry data sets is cellular sub-type classification. Clustering is one of the most challenging steps as it forms a basis for all subsequent differential tests on marker expression for biomarker discovery and population abundance analysis. Identification of cell populations depends on the number of features measured and cytometry has been able to push the detection limit to over 50 parameters per cell. However, increased dimensionality makes it difficult to capture the underlying heterogeneity of the data. Dimensionality reduction methods maximize the variance in the data and reduce the number of variables by mapping it onto a lower-dimensional space. Principal component analysis (PCA), a linear dimensionality reduction method, represents the original data in 2 or 3 dimensions by using a linear combination of the original feature vectors and maps data points onto orthogonal dimensions, which explain the maximum variance. However, PCA fails to capture the non-linearity and asynchronous nature of single-cell data, which is better visualized using non-linear dimensionality reduction techniques like t-SNE or uniform manifold approximation and projection or UMAP [16] (**Box 1**). t-SNE represents each cell in a lower dimensional manifold that is computed using the Barnes-Hut implementation of the t-stochastic neighbor embedding (t-SNE) algorithm [17]. t-SNE is currently one of the most popular methods of representing single-cell data.

Briefly, t-SNE computes a pairwise similarity matrix between all cells using a distance metric calculated from the feature vectors in high dimensions. Next, it initializes each cell to a random starting location in the 2 or 3 t-SNE dimensions and computes a second lower dimensional similarity matrix. The algorithm tries to iteratively minimize the difference between the lower and higher dimensional similarity matrices thereby updating the location of each cell in 2 or 3 dimensions. t-SNE optimization follows a stochastic nature, therefore every compilation of the method leads to slightly different lower manifold projections. It is thus advisable to run the method multiple times in order to achieve a concise representation of the variability of the different results. Furthermore, cells that are alike in higher dimensions are usually clustered together in t-SNE space. However, the opposite is not always true and thus this warrants caution with the analysis of t-SNE plots.

In order to explore the features of a t-SNE analysis, we applied the Rtsne package in R on an original CyTOF data-set [18], which measures more than 40 surface markers in a healthy human CD4^+^ T cell sample from peripheral blood mononuclear cells (PBMC). Our goal was to substantiate whether t-SNE was capable of recovering the major T-cell subtypes using a combination of surface markers and secreted cytokines. t-SNE cannot conveniently process very large data-sets and suffers from slow computation time. Since cytometry allows high-resolution dissection of cellular parameters, it usually measures a much larger number of cells, which can significantly increase time complexities of the analyses. Down-sampling the data set usually overcomes such complexities. Density-dependent down-sampling detects regions of density within the data-set and then downsizes keeping the structure and distribution consistent. This is beneficial for rare cell populations to define their own clusters instead of being subsumed under the highly abundant cell types. We performed density-dependent down-sampling of the PBMC CyTOF dataset using the SPADE [10] package. The algorithm was run using lineage markers and default parameter settings. We observed that the surface markers could delineate most of the different subsets. We could define the different memory subsets within PBMC using marker profiles thereby providing information about the potential for cells to home and migrate (**Figure 2A**). Moreover, overlapping clusters suggest that CD4^+^ T cells are plastic and display the ability to differentiate from one to another subtype. Different sequential t-SNE runs using identical parameters resulted in slightly different results due to differing events sampling conditions. t-SNE has been implemented in CRAN (http://www.r-project.org/) and it is also available as a plugin in FlowJo as well as in Cytobank at *www.cytobank.org*.

**Figure 2.**
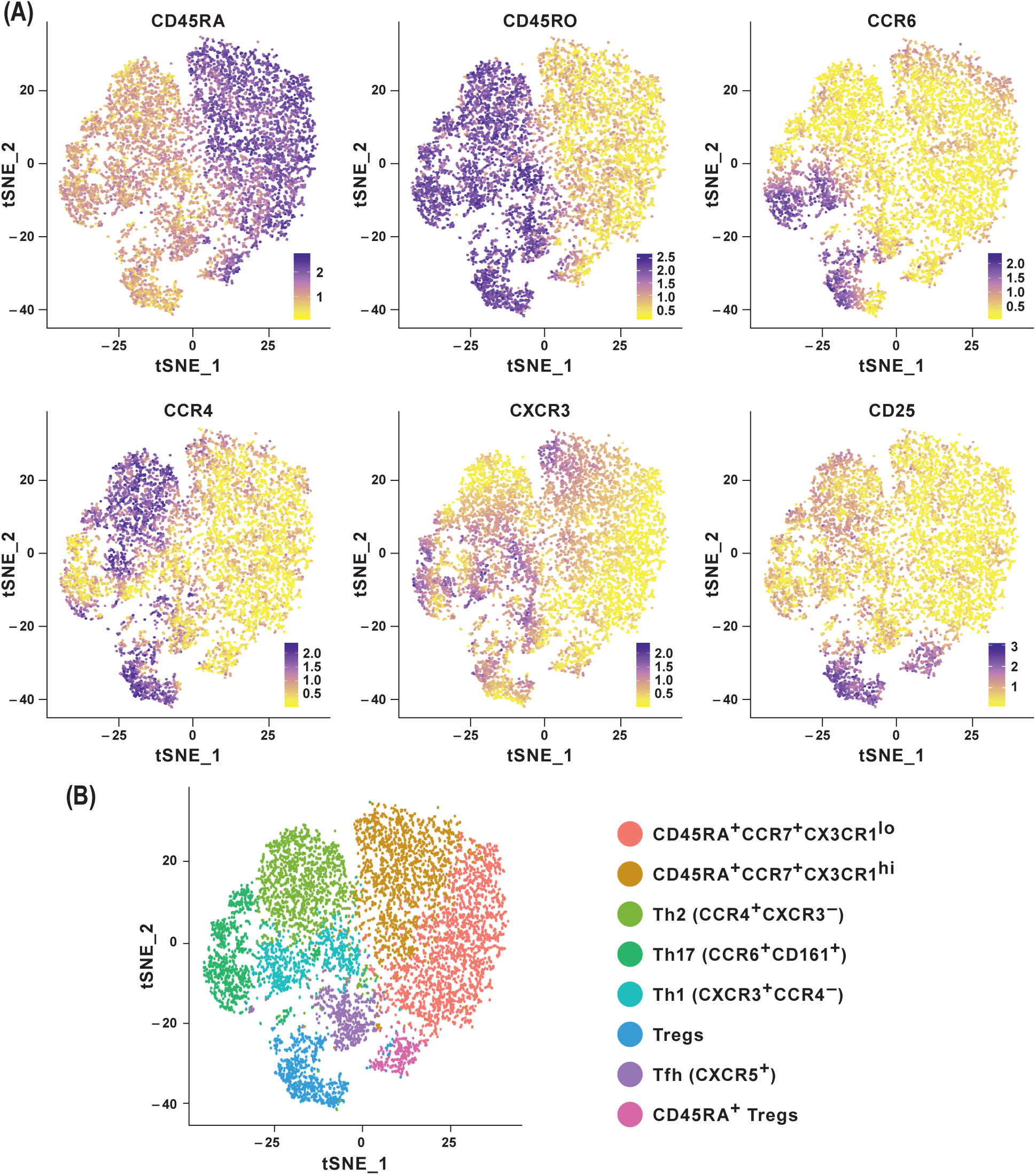
Dimensionality reduction using tSNE shows cellular heterogeneity. **(A)** tSNE plots for CD4^+^ PBMC from publicly available original CyTOF data (Wong et al. *Immunity* 2016) colored by selected marker intensities (median expression). **(B)** Identification of major immune subsets defined on the basis of lineage marker expressions from (A). Markers selected for tSNE analyses should represent proteins that delineate classic cellular lineages and differentiation states. Multiple tSNE runs can lead to slightly different results due to minor differences in computations. Despite this situation, multiple runs on the same dataset will produce highly similar results and the takeaway message will essentially be the same.

Diffusion maps, similar to PCA, are mainly used for data visualization in a non-linear fashion and can be a classic tool to investigate continuity in single-cell data. They were introduced by Coifman et al. [24] and are constructed using eigenvalue decomposition of a random-walk based transition matrix, which was recently adapted in Haghverdi et al. [9] to the single-cell setting. The method preserves the global relations between cells and has been able to successfully capture the developmental trajectories of differentiating cells, along with branching events, enabling it to capture both abundant as well as rare cell populations.

### Box 1

Non-linear dimensionality reduction methods scaling better than t-SNE

A recently introduced analysis technique for high-dimensional cytometry data known as HSNE or Hierarchical Stochastic Neighbor Embedding transcends the scalability limit of conventional t-SNE [19]. HSNE constructs a hierarchy of non-linear similarities between events that can be explored interactively up to single-cell details. The utility of this technique is to identify rare cell types that might otherwise be missed during down-sampling. HSNE has been implemented by *Unen et al.* as an integrated analysis tool Cytosplore [20].

Another non-linear dimensionality reduction technique which has garnered special is UMAP [16]. It is based on a novel manifold learning technique and can potentially better preserve the global structure of the data compared to t-SNE along with preserving local neighborhood aspects. Also, it scales better than most t-SNE packages in embedding large high-dimensional datasets which makes them a viable choice as a general purpose dimensionality reduction method.

## Organizing Single cells as Clusters for Sub-type Classification

Many tools now exist that group cells into discrete sub-populations based on feature space such as SPADE, FlowSOM etc. and employ unsupervised techniques for visualization of high-dimensional cytometry data. SPADE or spanning-tree progression analysis of density-normalized events organizes cellular populations into hierarchies based on similar phenotypes [10]. It provides an intuitive 2D depiction of multiple cell-types in a branched tree structure (**Box 2**). A typical SPADE tree is comprised of nodes representing cell clusters, which are further connected through edges, which represent relationships and provide information about the underlying similarity of cell-types [21]. Only the connections between nodes via edges can be used to draw conclusions about cluster similarities. The larger the distance between two connected clusters, the more dissimilar are the features of the events within those clusters. Additionally, a SPADE tree can be colored using the expression level of any preferred marker giving insights into the differential expression pattern between the events from different clusters.

### Box 2

Hierarchical tree representation of single-cells using SPADE

Typically, SPADE begins by performing a density-dependent down-sampling of the raw dataset followed by an unsupervised agglomerative hierarchical clustering to identify distinct sub-populations. It then builds a minimum spanning tree representation to link the clusters beginning with a randomly chosen but already connected subgraph and adding an edge to it iteratively. Finally, it performs up-sampling by assigning all cells in the initial dataset to the clusters identified [21].

Ideally, SPADE can recover cellular hierarchy corresponding to known biology from high-dimensional cytometry data-sets. However, performance is limited by a number of user-defined factors such as the desired number of clusters, outlier density and target density following down-sampling which can affect the detection of rare cells. Furthermore, since SPADE is a non-deterministic method and the minimum-operation in the spanning-tree step is sensitive with respect to outliers, every run would result in a distinctly different tree structure.

We analyzed the performance of SPADE using the CD4^+^ T cell CyTOF data set from PBMC [18]. The data was transformed using the hyperbolic arcsine function. SPADE was applied to this dataset using all the surface markers and default parameter settings except for the number of clusters, which was set at 100 to better capture the heterogeneity of the data. To explore the underlying structure and heterogeneity in the data the SPADE trees were annotated using median expression of different markers (**Figure 3C**). The median marker intensities for CD45RA and CD45RO clearly indicate the presence of a naïve and memory T cell compartment in the peripheral blood. Additionally, by comparing expression of markers CD25 and CD127 one could also identify and delineate regulatory T cells from naïve and central memory T cells and effector T cells. SPADE is implemented as an R package and is also available from Cytobank.

**Figure 3.**
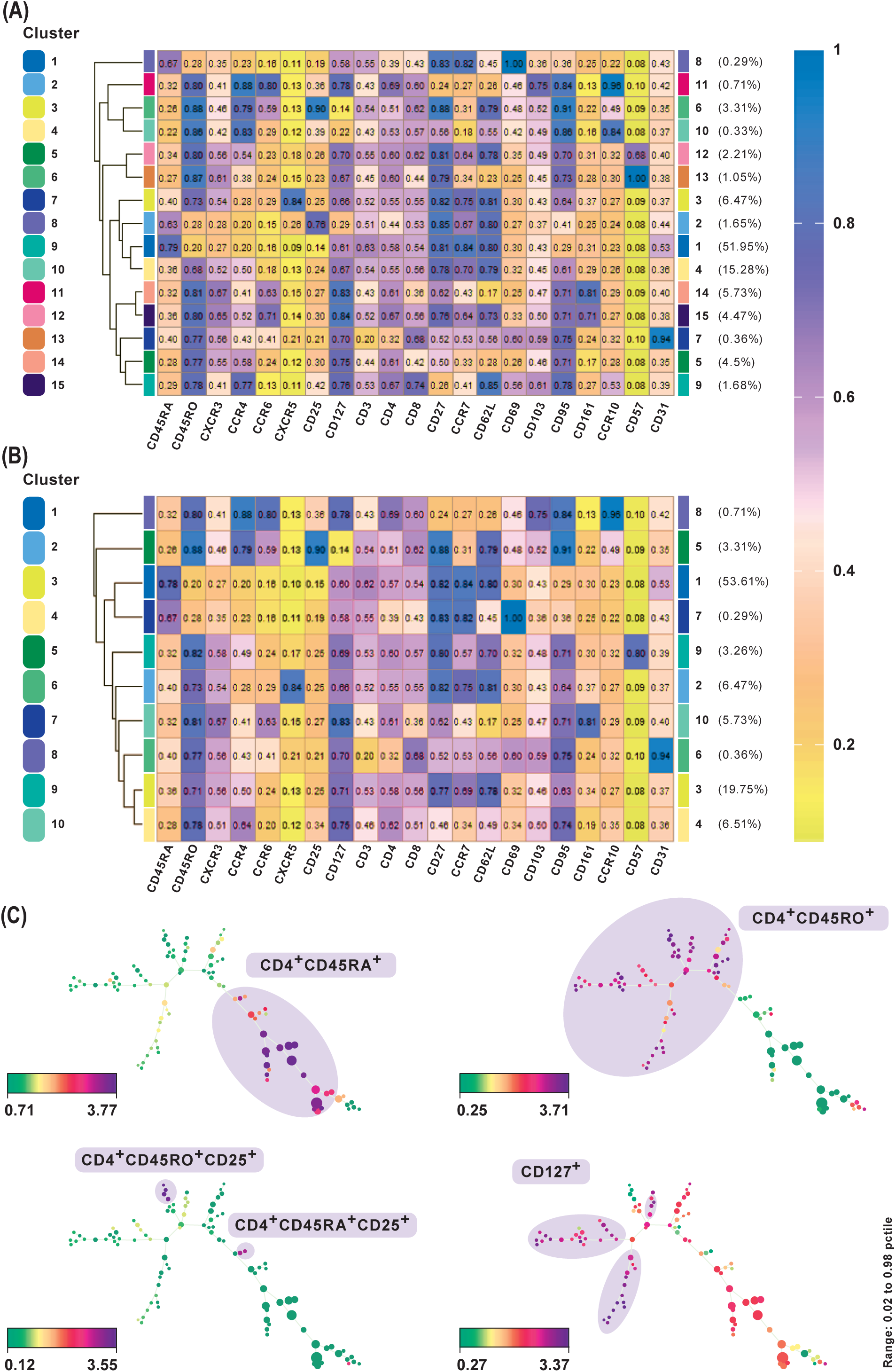
Subset classification of single cells by unsupervised clustering using SPADE and FlowSOM. **(A)** FlowSOM clustering with publicly available original CyTOF data from CD4^+^ human PBMC (*Wong et al. Immunity* 2016). Data was transformed using logicle transformation. Median intensities of 21 markers were used for analysis. FlowSOM provides a concise representation of the potential number of cell types present in the study. It can be used as a starting point for data exploration prior to biaxial gating. Meta-clustering with 15 clusters was able to identify the expected clusters associated with naive (CD45RA^+^), memory (CD45RO^+^), Th1 (CXCR3^+^CCR4^−^), Th2 (CXCR3^−^CCR4^+^), Treg cells, follicular helper T cells (CXCR5^+^) and Th17 cells (CCR6^+^CD161^+^). Additionally, by not down-sampling the raw data, it could potentially identify low frequency clusters as well, such as CD57^+^ and CD31^+^cells, which gives FlowSOM an advantage in being able to capture subtle differences between clusters based on their differential marker profiles. **(B)** One could also perform an expert-based merging of algorithm identified clustering. **(C)** SPADE analysis performed on the same data. Nodes in SPADE trees were overlaid with CD45RA, CD45RO, CD25 and CD127 marker expression for characterization of naïve (CD45RA^+^), memory (CD45RO^+^), as the two main branches, and regulatory T cell or Tregs (CD25^hi^CD127^lo^) populations. Multiple SPADE runs result in slightly differing trees, however, the cellular subtypes identified between runs with the same dataset are similar. The tree structure, as well as connectivity via edges, also give information regarding the degree of similarity between cell populations.

A central challenge in visualizing larger datasets is to achieve and maintain performance without compromising on speed. In line with this, Van Gassen *et al.* [11] introduced FlowSOM, which uses self-organizing maps (**Box 3**). In contrast to t-SNE and SPADE analyses, several plots are not required to determine an accurate cell-type classification of clusters and their boundaries.

We present results of FlowSOM clustering, which was applied to an original CD4^+^ PBMC CyTOF data set and expected to identify the known cell populations in the study [18]. The data was transformed using logicle transformation and scaled. FlowSOM was applied using the standard parameter settings and lineage markers for clustering, which we considered could positively delineate subsets. Notably, the method was able to detect both high as well as low frequency cell populations (**Figure 3A**). Meta-clustering with 15 clusters was able to identify the expected clusters associated with naive (CD45RA^+^), memory (CD45RO^+^), Th1 (CXCR3^+^CCR4^−^), Th2 (CXCR3CCR4^+^), Treg cells, follicular helper T cells (CXCR5^+^) and Th17 (CCR6^+^CD161^+^). By averting down-sampling, it could potentially identify low frequency clusters as well, such as CD57^+^ and CD31^+^, giving FlowSOM an advantage in being able to capture subtle differences between clusters based on their differential marker profiles.

### Box 3

#### Self-organizing maps of single-cells

Self-organizing maps (SOMs) are a classical unsupervised dimensionality reduction and clustering technique, which trains a map on a discretized representation of the input space to produce a low-dimensional embedding of the same. SOMs are designed to preserve the topological information of the original input space by mapping similar high-dimensional data points to the same region in 2D space [22].

SOMs in FlowSOM are trained on the input matrix which performs the embedding of the higher dimensional space onto a rectangular grid. The resulting grid of nodes already correspond to cells, clustered using nearest neighbors, and can be visualized as star charts using mean marker expressions. The nodes of the self-organizing map are connected in a minimum-spanning tree for graphical representation, providing results comparable to SPADE visualizations. An additional meta-clustering step is performed, which includes a much larger number of nodes than clusters to give a detailed overview of the data with subtle differences. FlowSOM can be used alongside a manual gating analysis to easily compare the results in addition to examining events which are traditionally “gated out”. FlowSOM clustering does not mandate subsetting of the data because it scales easily and does not require down-sampling [23]. Thus, by visualizing all cells simultaneously, annotating cell types become easier and the risk to lose novel cell populations is reduced.

### Trajectory Inference of Differentiating Cells and Graph Abstraction

Cellular differentiation is a non-linear and continuous phenomenon [4]. Additional information can be gained from aligning asynchronously differentiating cells according to their inherent developmental state (**Figure 4D**). Their temporal order can be computed from expression profiles and measured using a random-walk-based distance metric known as diffusion pseudo time (DPT) [14] (**Box 4**). DPT identifies developmental progression, branching points as well as differential expression of key decision-making cell biomarkers on the single-cell level. DPT has been used to analyze an InDrop single-cell RNAseq data from Klein et al [25], where it revealed differentiation and transcription factor dynamics of mouse embryonic stem cells after leukemia inhibitory factor withdrawal and identified major clusters with different biological functions. Notably, the analysis identified one cluster enriched for pluripotency factors that were active during early pseudo-time [14].

**Figure 4.**
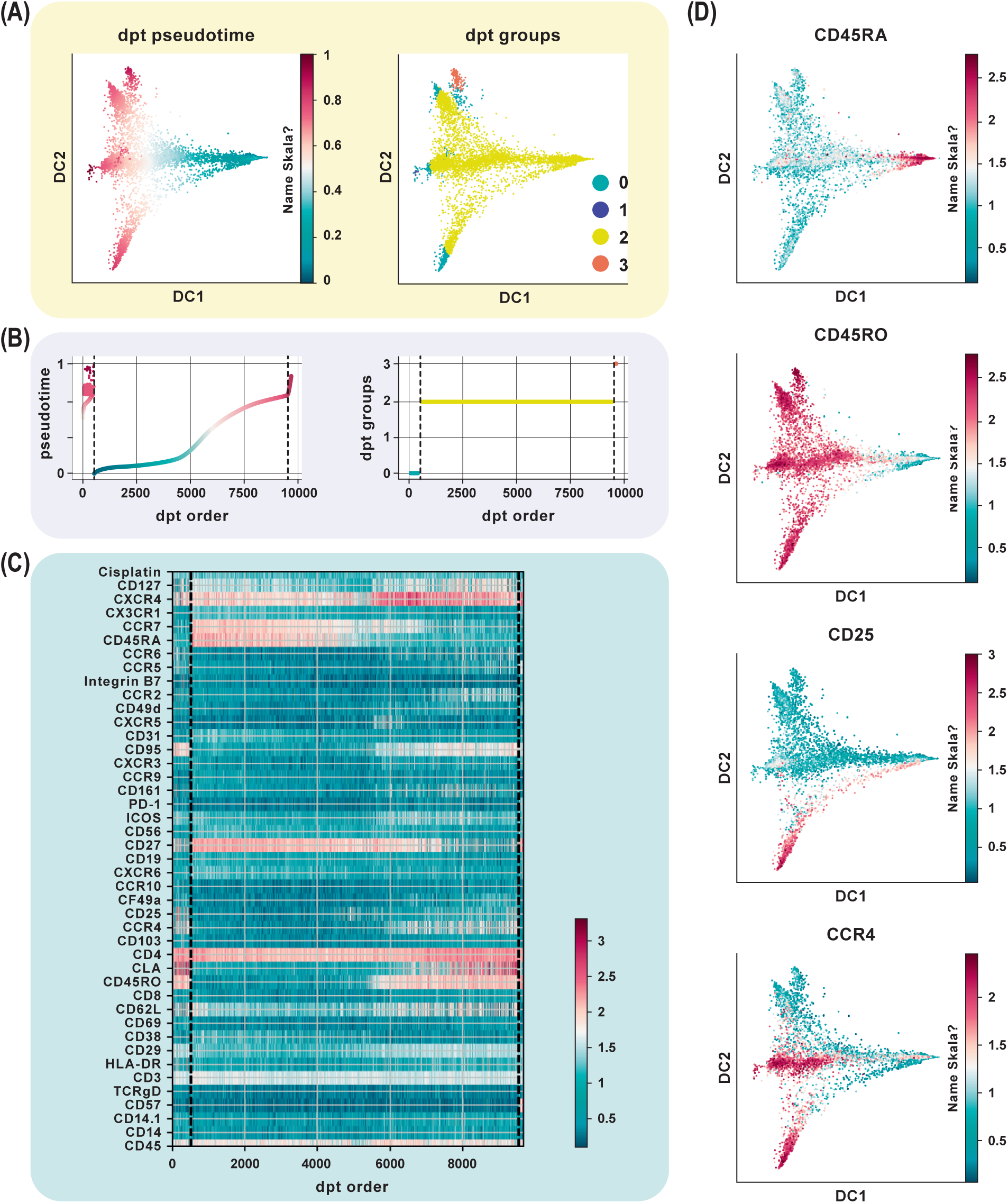
Trajectory inference from static snapshot cytometry data. **(a)** Application of DPT to CD4^+^ PBMC using an original CyTOF dataset (Wong et al. *Immunity* 2016). Clustering based on the tip cells identifies three major DPT groups. The root cell, with a high expression for CD45RA (naïve compartment), has the lowest pseudotime. **(B)** The three DPT groups and their corresponding change in pseudotime. **(C)** Heat-map of marker expression with cells ordered by DPT and depicts protein marker dynamics with cells transitioning from CD45RA^+^ to CD45RO^+^ as seen in DPT group 2. **(D)** Diffusion maps of CyTOF data colored by selected lineage marker intensity profiles, which identify major immune subsets aligned along internal developmental states of the data. The ensemble of diffusion plots clearly reveal the major T cell subsets, e.g. cells differentiating from a naïve to a memory state with Treg cells, T helper subsets such as Th1 and Th2 cells populating the ends of the trajectory.

This method reveals new biology from a variety of experimental settings through robust computation of pseudo-time and scalability. In comparison to previous algorithms for pseudotemporal ordering, such as Wanderlust/Wishbone [13,26] and Monocle [12], DPT’s random-walk-based formulation has been shown to perform significantly better in ordering cells according to pseudotime. Unlike DPT, Monocle utilizes an only partially robust minimum spanning tree approach and is unable to scale to high cell numbers. Wishbone, on the other hand, computes pseudotime distance based on shortest paths on graphs, which leads to a complicated and iterative computation to account for branches [6]. Since then many more algorithms for trajectory inference have been proposed, especially for scRNAseq. Saelens *et al.* [27] perform an extensive and comprehensive assessment of 29 published trajectory inference methods on both simulated as well as real datasets and provide a set of guidelines for users.

#### Box 4

##### Inferring cellular trajectories using diffusion pseudotime

The DPT algorithm first computes a transition matrix from single-cell expression data by convolving Gaussian kernels centered at nearby cells, effectively constructing a weighted nearest-neighbor graph of the data. Next, it determines the probabilities for each cell to transition to each other cell in the data set using random walks of any length on this graph. These transition probabilities correspond to edge weights. These walks can be considered as a proxy for the cells’ probabilities of differentiating toward different cell types. The probabilities for each cell are stored in a vector, and the DPT between two cells is calculated as the Euclidean distance between their two vectors. The developmental progression of each cell in the data set is then measured by computing its DPT with respect to a specified root cell [14].

We evaluated the performance of diffusion maps and DPT on the CD4^+^ T cell CyTOF dataset after following a density dependent down-sampling and using logicle transformation [18]. The root cell was chosen as the cell having the minimum expression for CD45RO, based on the knowledge that naïve T cells express CD45RA and that upon antigen exposure they differentiate into central and effector memory T cells gaining expression of CD45RO and losing expression of CD45RA (**Figure 4A**). The ensemble of diffusion plots clearly revealed the major T cell subsets, e.g. transition from naïve to memory T cells with Treg cells and T helper subsets originating towards the end of differentiation (**Figure 4D**). The heat-map orders cells by DPT and depicts protein marker dynamics with cells transitioning from CD45RA^+^ to CD45RO^+^ (**Figure 4C**).

Many unsupervised single-cell data analysis algorithms are based on clustering approaches which label groups of cells into discrete clusters with biologically distinct phenotypic and functional characteristics. On the other hand, trajectory inference algorithms assume that data lie on a connected manifold and project cells on a so-called pseudotime by computing paths between them using some distance metric along this manifold. Partition-based graph abstraction (PAGA) combines analysis strategies of both clustering as well as trajectory modeling to compute an abstracted graph representing the overall topology of a possibly disconnected manifold of cells [15]. It first computes a neighborhood graph of single cells whose partitions represent groups of similar cells. From this, it generates a simple abstracted graph whose nodes correspond to these partitions and edges represent a confidence measure for the connectivity between partitions. The method utilizes a random-walk-based distance measure to generate a topology preserving map of the underlying differentiation manifold from single-cell measurements and is shown to be computationally efficient [15].

We applied PAGA on the CD4^+^ T cell PBMC CyTOF dataset using default settings and a resolution of 0.6 (**Figure 5A**) [18]. The abstracted graph was able to reconstruct the major T cell subsets arising from CD4^+^ T helper cell differentiation. Cluster 1, 2, 3 and 4 constituted the memory compartment while cluster 0 and 5 represent naïve subsets. Cluster phenotypes could be identified from the marker expression distribution (**Figure 5B**). In datasets with inherent continuous manifolds, PAGA constructs a tree-like lineage graph with disconnected clusters. Together it explains the global topology of the data and also reconstructs differentiation processes.

**Figure 5.**
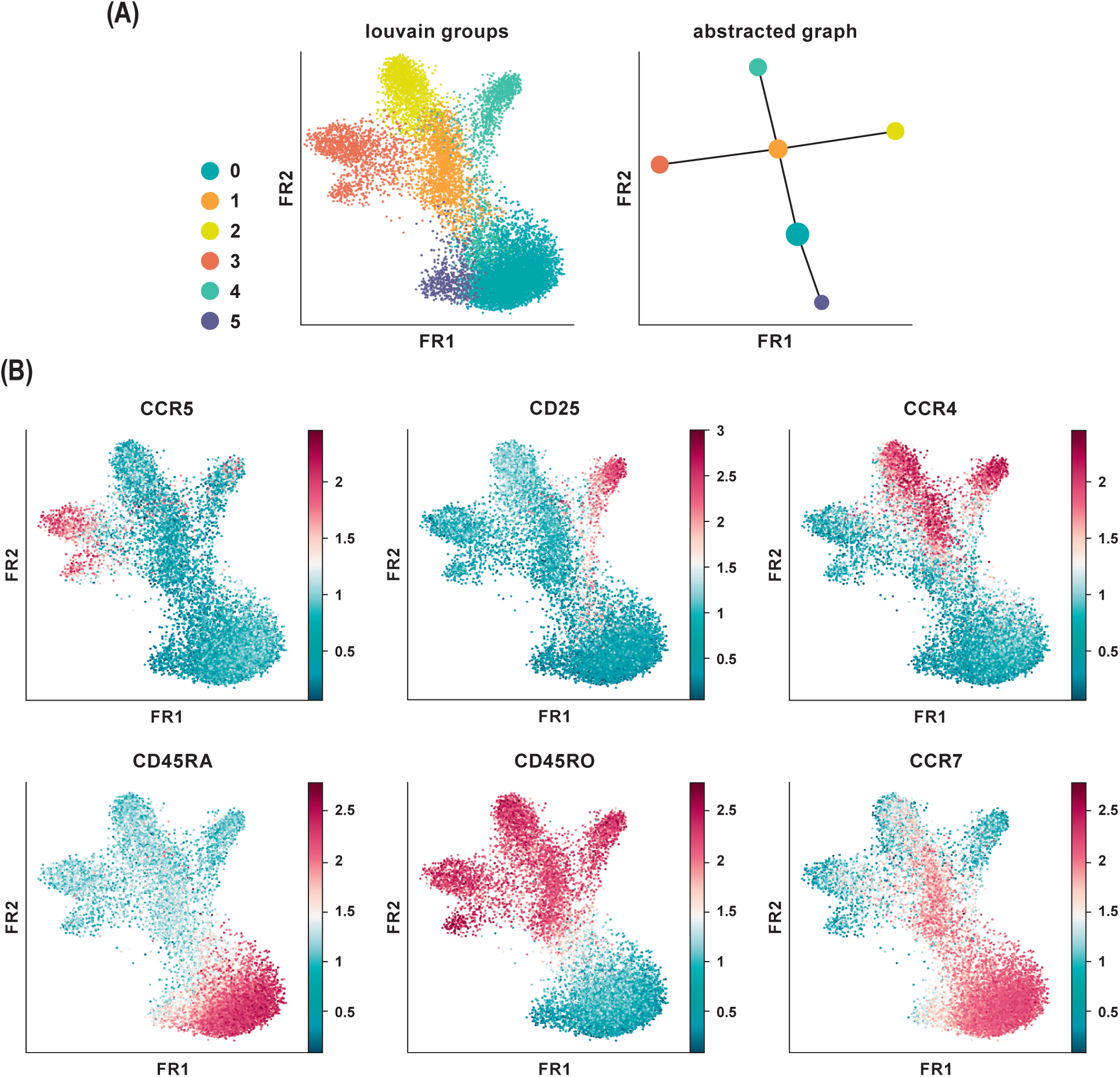
Partition-based graph abstraction of CyTOF data. **(A)** Shown is a data analysis of an original CyTOF data-set depicting CD4^+^ T cells derived from PBMC (Wong et al. *Immunity* 2016). Louvain groups in a single-cell graph visualized using the Fruchtman-Reingold (FR) algorithm, which conserves continuous structure in data better than tSNE. Abstracted graph visualized using a simple tree based graph drawing layout. **(B)** Identification of cluster phenotypes from marker expression distribution. In data-sets with inherent continuous manifolds, PAGA constructs a tree-like lineage graph with disconnected clusters. Together, it explains the global topology of the data as well as reconstructs differentiation processes.

### Concluding Remarks and Future Perspectives

Single-cell technologies have become a ubiquitous tool as researchers realize their untapped potential to uncover cellular heterogeneity and functionality at a greater resolution than with bulk analysis. The past few years have seen a significant change in the ways cytometry datasets are being analyzed and a wealth of novel computational tools is now available to mine complex and high-dimensional data in an unbiased automated manner. Depending on the biological question, one of several computational methods can be incorporated to potentially substantiate the findings of manual gating as well as deepen our understanding of how the immune system functions in health and disease.

Many integrated data analysis frameworks now exist to facilitate a comprehensive interrogation of high-dimensional single-cell data. Most of these have been originally developed for scRNA-seq data. However, they can be extended for the purpose of cytometry as well. Cytofkit, an integrated analysis pipeline, is specially designed for mass cytometry data and is available as a Bioconductor package [30]. It also provides a graphical user interface as well as a Shiny application for interactive and effortless usage and visualization of results. Seurat, also available as a Bioconductor package, contains implementations of commonly applied analytical techniques for exploring single-cell expression data [31]. Scanpy, a Python frame-work, provides computationally efficient and state-of-the-art methods to address the statistical challenges associated with scRNA-seq data [32]. All of these packages incorporate both novel as well as established methods to perform data pre-processing, feature selection, linear and non-linear dimensionality reduction, standard unsupervised clustering algorithms for automatic detection of cell subsets and differential testing. Scanpy additionally integrates novel in-house algorithms and performs trajectory inference. One of the most productive research areas in future should be towards developing and maintaining such integrated analysis pipelines for cytometry data as well as bridging the gap with other OMICS data analysis for a more comprehensive interpretation of study models.

There are many more established algorithms for single-cell analysis mostly developed for scRNA-seq that also allow investigation of cytometry datasets, several of which have been reviewed earlier [6,33,34]; for a large-scale overview please consider this list [35]. However, we find that the different methods vary significantly in terms of scalability, speed and computational skill required to interpret results. We discuss and demonstrate the feasibility and power of several current computational tools to translate complex static snapshot data obtained from high-dimensional single-cell datasets into dynamic ontological and regulatory networks of the immune system. A potential avenue for further development would be to incorporate machine learning methods to infer developmental trajectories directly from cytometry data, which currently describes much less features to model the underlying manifold and is, thus, a limitation. Ultimately, it depends on the experience and requirement of the investigator to make an informed decision about the choice of the data exploratory method.

## Acknowledgements

Funding: This work was supported by the German Research Foundation (SFB1054 Teilprojekt B10 to C.E.Z, SFB1335 Teilprojekt P18 to C.E.Z.), the Fritz-Thyssen Stiftung (C.E.Z), and the German Center for Infection Research (C.E.Z.).

